# An engineered xCas12i with high activity, high specificity and broad PAM range

**DOI:** 10.1101/2022.06.15.496255

**Authors:** Hainan Zhang, Xiangfeng Kong, Mingxing Xue, Zikang Wang, Yinghui Wei, Haoqiang Wang, Jingxing Zhou, Weihong Zhang, Mengqiu Xu, Xiaowen Shen, Jinhui Li, Jing Hu, Na Zhong, Yingsi Zhou, Hui Yang

## Abstract

The type-V CRISPR effector Cas12i, with its smaller size, short crRNA guiding, and self-processing features, is a potentially versatile genome editing tool. By screening Cas12i proteins from a metagenomic database, we identified a natural variant with high activity in mammalian cells, named as xCas12i. We further engineered the PAM-interacting, REC, and RuvC domains for enhanced cleavage activity and specificity. This variant, named as high-fidelity Cas12Max, exhibited robust genome editing activity and minimal off-target activity with a broad 5’-TN recognition profile. With the fusion of deaminase TadA8e and further optimization of xCas12i, the base editor dCas12i-Tad8e also showed the high editing efficiency. This study provides highly efficient and specific tools for gene therapy.

## Introduction

The clustered regularly interspaced short palindromic repeats-Cas (CRISPR-Cas) systems, including type II Cas9 and type V Cas12 systems, which serve in the adaptive immunity of prokaryotes against viruses, have been developed into genome editing tools^1-3^. Compared with type II systems, the type V systems including V-A to V-K showed more functional diversity^4, 5^. Amongst them, Cas12i has a relatively smaller size (1033-1093 aa), compared to SpCas9 and Cas12a, and has a 5’-TTN protospacer adjacent motif (PAM) preference^4, 6, 7^. Cas12i is characterized by the capability of autonomously processing precursor crRNA (pre-crRNA) to form short mature crRNA. Cas12i mediates cleavage of dsDNA with a single RuvC domain, by preferentially nicking the non-target strand and then cutting the target strand^8-10^. These intrinsic features of Cas12i enable multiplex high-fidelity genome editing. However, the natural variants of Cas12i (Cas12i1 and Cas12i2), showed low editing efficiency which limits their utility for therapeutic gene editing.

To address these limitations, we screened ten natural Cas12i variants and found one, xCas12i, with robust high activity in HEK293T cells. Engineering of xCas12i by arginine substitutions at the PAM-interacting (PI), REC and RuvC domains led to the production of a variant, high-fidelity Cas12Max (hfCas12Max), with significantly elevated editing activity and minimal off-target cleavage efficiency. In addition, we assessed the base editing efficiency of xCas12i-based base editor, and thus expanded the genome-editing toolbox.

## Results

### Identification and characterization of type V-I systems xCas12i

In order to identify more Cas12i variants, we developed and employed a bioinformatics pipeline to annotate Cas12i proteins, CRISPR arrays and predicted PAM preferences, and found 10 new CRISPR/Cas12i systems. To evaluate the activity of these Cas12i variants in mammalian cells, we designed a fluorescent reporter system which detected the increased enhanced green fluorescent protein (EGFP) signal intensity activated by Cas-mediated dsDNA cleavage or double strand breaks (Fig.S1a). This system relied on the co-transfection of a plasmid coding for mCherry, a nuclear localization signal (NLS) -tagged Cas protein and its guide RNA (gRNA) or crRNA, and one coding BFP and activatable EGxxFP cassette, which is EGxx-target site-xxFP^11^. EGFP activation was carried out by Cas mediated DSB and single-strand annealing (SSA)-mediated repair. Using this system, we observed that a variant, xCas12i, with targeted crRNA induced significant activation of EGFP expression (Fig. 1a, Fig. S1b), and exhibited a higher editing frequency than LbCas12a or SpCas9 as determined by Fluorescence Activated Cell Sorter (FACS) analysis (Fig. 1a). The xCas12i variant was smaller in size compared to SpCas9 and LbCas12a (Fig. S2a). We explored the effects of spacer length on cleavage efficiency in xCas12i, and found that 17-22 nt were optimal length for their activation (Fig. S2b). Considering the 5’-TTN PAM preference of Cas12i, we performed a NTTN PAM identification assay using the reporter system. We found that xCas12i showed a consistent high frequency of EGFP activation at target sites with 5’-NTTN PAM sequences, while LbCas12a had comparable activity at 5’-TTTN PAM, respectively (Fig. S2c).

**Fig.1.**
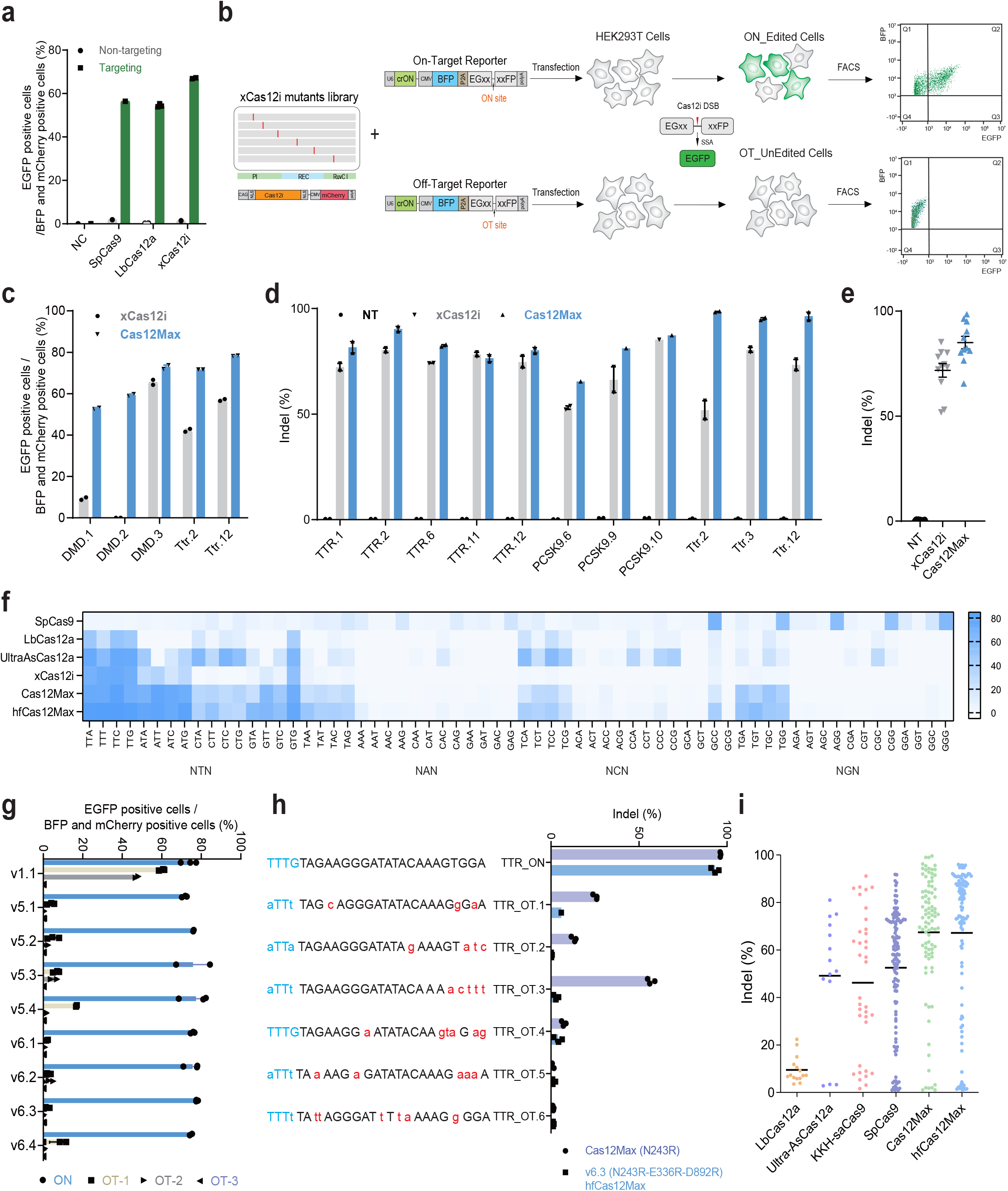
hfCas12Max, an engineered natural variant xCas12i, mediated high-efficient and -specificity genome editing in mammalian cells. **a**, A natural variant xCas12i mediated EGFP activation efficiency determined by flow cytometry. **b**, Schematics of protein engineering strategy for mutants with high efficiency and high fidelity using an activatable EGFP reporter screening system with on-targeted and off-targeted crRNA. **c-e**, Cas12Max exhibited significantly increased cleavage activity than xCas12i at various genomic or reporter plasmids target sites. **f**, Both Cas12Max and hfCas12Max exhibited a broader PAM recognition profile than other Cas proteins, including 5’-TN and 5’-TNN PAM. **g**, v6.3 reduced off-target at OT.1 and OT.2 sites and retained indel activity at TTR-ON targets, compared to v1.1-Cas12Max. **h**, TIDE analysis showed that v6.3-hfCas12Max retained comparable activity at TTR.2-ON targets and almost no at 6 OT sites, to Cas12Max. **i**, Comparison of indel activity from Cas12Max, hfCas12Max, LbCas12a, Ultra AsCas12a, SpCas9 and KKH-saCas9 at TTR locus. hfCas12Max retained the comparable activity of Cas12Max, and higher gene-editing efficiency than other Cas proteins. Each dot represents one of three repeats of single target site.

To further confirm the dsDNA cleavage activity of xCas12i in mammalian cells, we transfected an all-in-one plasmid encoding NLS tagged xCas12i with crRNAs targeting 37 sites from *TTR*^*12*^, *PCSK9*^*13*^ in HEK293T or *Ttr* in N2a cells. The editing efficiency, *i*.*e*. indel (insertion and deletion) formation at these loci was measured 48 hours after transfection using FACS and targeted deep sequencing (Fig. S2d). We found that xCas12i mediated a high frequency, up to 90%, of indel formation at most sites from *Ttr, TTR* and *PCSK9*, with a mean indel formation rate of over 50% (Fig. S2e-f). These data indicate that xCas12i exhibits a robust genome editing efficiency in mammalian cells, suggesting it has excellent potential for therapeutic genome editing applications.

### Engineered xCas12i mediates high-efficiency editing at 5’-TN PAM sites

To enhance its activity and expand its scope of PAM site recognition, we sought to engineer xCas12i protein via mutagenesis and screen for variants with higher efficiency and broader PAM using a reporter system, similar to what is described above. Substitution of an amino acid in the DNA-binding pocket with positively charged arginine (R) was shown to enhance the activity of the type V system^14-16^. Combined with predictive structural analysis of xCas12i, we performed an arginine scanning mutagenesis approach in the PI, REC-I and RuvC-II domains, generating a library of over 500 mutants (Fig. 1b, Fig. S3a). We then individually transfected these mutant variants with an activatable EGFP reporter system in HEK293T cells and analyzed them by FACS (Fig. 1b). Based on the fluorescence intensity of cells with activated EGFP, over 100 mutants showed an increased frequency of activated cells relative to wild type (WT) xCas12i, and one mutant, named as Cas12Max, containing N243R showed a 3.4-fold improvement (Fig. S3a). We next targeted *DMD* or *Ttr* sites using the fluorescent reporter system, and found that Cas12Max displayed a markedly increased frequency of EGFP activation, relative to WT xCas12i (Fig. 1c, Fig. S4a-b). To further test the efficacy of Cas12Max in targeting genomic loci, we designed a total of eight gRNAs to target sites *TTR* and *PCSK9* in HEK293T cells and three more targeting *Ttr* in N2a cells. Consistent with our previous result, Cas12Max exhibited a significantly increased frequency of indels compared to WT xCas12i (Fig. 1d-e).

Additionally, to investigate Cas12Max’s PAM preference, we performed a 5’-NNN PAM recognition assay by designing reporter plasmids with the same target sequence but different PAM. Besides showing a consistent or higher cleavage activity at sites with a 5’-TTN PAM, Cas12Max showed a similarly high cleavage activity for targets with TNN, ATN, GTN and CTN PAM sites, compared with the commonly used Cas12^7, 17^ (LbCas12a, Ultra-AsCas12a) (Fig. 1f). Taken together, these results demonstrate that Cas12Max exhibits high-efficiency editing activity with highly flexible 5’-TN or 5’-TNN PAM recognition.

### hfCas12Max mediates high-efficient and specific genome editing

To examine the specificity of Cas12Max, we transfected a construct designed to express it with crRNA targeting *TTR*^*12*^, and performed indel frequency analysis of on- and off-target (OT) sites predicted by Cas-OFFinder^18^. Using reporter system or targeted deep sequence analysis, we found that Cas12Max efficiently edited target sites and resulted in significant indel formation at 2 of 3 predicted off-target sites (Fig. S5). To eliminate the off-target activity of Cas12Max, we screened these mutants with mutations in the REC and RuvC domains^19^, which have undiminished on-target cleavage activity, for those with no off-target activity, using two activatable reporter systems, each containing one OT site (Fig. 1b). We found that four mutants (v4.1-V880R, v4.2-M923R, v4.3-D892R and v4.4-G883R) maintained a high level of on-target editing activity and showed significantly reduced off-target EGFP activation (Fig. S6a). We further combined these four amino acid substitutions with N243R and/or E336R of Cas12Max and found that the variant v6.3 (N243R/D892R/G883R) showed the lowest off-target EGFP activation at OT.1 and OT.2 sites and high on-target at the ON.1 site (Fig. 1g, Fig. S6b-c). Targeted TIDE analysis of endogenous TTR.2 site and its off-target sites in HEK293T showed that v6.3 (N243R/D892R/G883R) significantly reduced off-target indel frequencies at six OT sites and retained on-target at ON site, compared to Cas12Max (Fig. 1h). In addition, relative to Cas12Max (v1.1), v6.3 (N243R/D892R/G883R) retained comparable or even higher on-target activity at DMD.1, DMD.2 and DMD.3 sties (Fig. S6d). Therefore, we named v6.3 as high-fidelity Cas12Max (hfCas12Max).

To comprehensively evaluate the performance of hfCas12Max in human cells, we designed large number of target sites in the exons of *TTR* for various Cas nucleases. In total, editing activity was monitored over 30 sites for hfCas12Max with TTN PAMs, 40 sites for SpCas9 with NGG PAMs, 4 sites for LbCas12a with TTTN PAMs, 4 sites for Ultra AsCas12a, and 7 sites for KKH-saCas9 with NNNRRT PAMs. Indel analysis showed that hfCas12Max exhibited an average efficiency of 70%, higher activity than other Cas nucleases, comparable activity with Cas12Max (Fig. 1i, Fig. S7a). Using 5’-NNN PAM reporter system, hfCas12Max maintained the broad 5’-TN and 5’-TNN PAM recognition profile similarly to Cas12Max (Fig. 1f). To further evaluate the specificity of hfCas12Max in human cells, we determined indel frequencies of P2RX5 and NLRC4 on-target and their corresponding *in silico* predicted off-target sites^20^. TIDE analysis showed that hfCas12Max had a higher on-target editing efficiency and similarly almost no indel activity at potential off target sites, compared to Ultra AsCas12a and LbCas12a (Fig. S8a-b). Overall, these results demonstrate that hfCas12Max has high efficiency and specificity and is superior to SpCas9 and other Cas12 nucleases.

### dxCas12i-based base editors efficiently mediate base editing in mammalian cells

We further explored the base editing of xCas12i by generating a nuclease-deactivation xCas12i (dxCas12i). This was done by first introducing single mutations (D650A, D700A, E875A, or D1049A) in the conserved active site of xCas12i based on alignment to Cas12i1^8^ and Cas12i2^10^ (Fig. S9a-b). Then, these variants were fused with TadA8e^V106W^ or human APOBEC3A (hA3A^W104A^) to form the dxCas12i base editors TadA8e.1-dxCas12i and hA3A.1-dxCas12i, respectively^21-23^. The initial versions of TadA8e.1-dxCas12i and hA3A.1-dxCas12i showed low base editing activity with frequencies of 8% A-to-G and 2% C-to-T, respectively (Fig. 2a-d). To address this, we introduced single and combined mutations for high cleavage activity into the PI and Rec domains of dxCas12i, which resulted in significantly increased A-to-G editing activity. Among the improved variants, dxCas12i-TadA8e-v2.2 (N243R/E336R) achieved 50% activity at A9 and A11 sites of the *KLF4* locus, markedly higher than the 30% activity of dLbCas12a-TadA8e (Fig. 2b, Fig. S10a-b). At target sites within *PCSK9*, TadA8e-dxCas12i-v2.2 showed a similarly increased efficiency to mediate A-to-G transitions, higher than dLbCas12a-TadA8e (Fig. S11). We then further engineered the NLS, linker, TadA8e protein and the N-terminus to produce dxCas12i-TadA8e-v4.3 which exhibited a nearly 80% A-to-G editing efficiency, while the editing activities of other dxCas12i-ABE versions were unchanged (Fig. 2c, Fig. S10c-d). In addition, the substitution of dxCas12i-v1.2 (N243R) or dxCas12i-v2.2 (N243R/E336R) showed consistently elevated C-to-T editing at C7 and C10 sites of *DYRK1A*, even higher at C7 by dLbCas12a-hA3A (Fig. 2d). These results together demonstrate that engineered dxCas12i-based editors exhibit the high base editing activity in mammalian cells.

**Fig.2.**
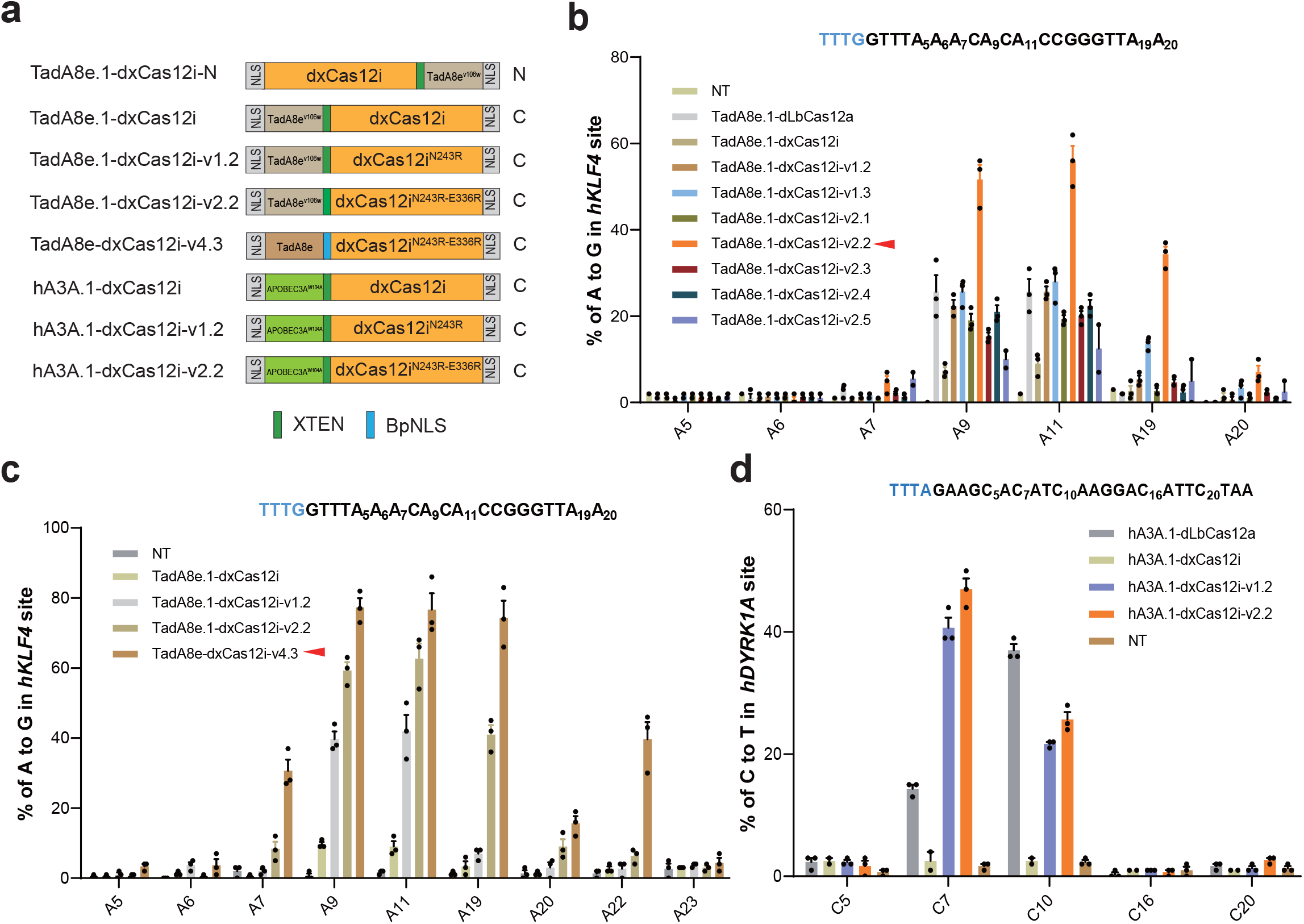
dCas12Max-TadA8e and dCas12Max-hA3A.1 mediated high base editing activity. **a**, Schematics of different versions of dCas12Max base editors. **b**, The A-to-G editing activity of TadA8e.1-dxCas12i-v2.2 was largely increased by combining v1.2 and v1.3 variant. **c**, Comparison of A-to-G editing frequency at the KLKF4 site from TadA8e.1-dxCas12i-v1.2, v2.2 and v4.3, ABE-dCas12Max (TadA8e-dxCas12i-v4.3) showed a high editing activity of 80%. **d**, Comparison of C-to-T editing frequency at the DYRK1A site from hA3A.1-dxCas12i, -v1.2 and v2.2, CBE-dCas12Max (hA3A.1-dxCas12i-v2.2) showed a high editing activity of 50%.

## Discussion

In this study, we demonstrate that the Type V-I Cas12i system enables versatile and efficient genome editing in mammalian cells. We found a natural Cas12i variant, xCas12i, that shows high editing efficiency at TTN-PAM sites. By semi-rational design and protein engineering of its PI, REC, RuvC domains, we obtained a high-efficiency, high-fidelity variant, hfCas12Max, which contains N243R, E336R, and D892R substitutions. In agreement with the hypothesis that introducing arginine at key sites could strengthen the binding between Cas and DNA, the introduction of N243R in the PI domain and E336R at REC domain significantly increased editing activity and expanded PAM recognition. Interestingly, D892R or G883R substitutions in the RuvC domain reduced off-target and retained on-target cleavage activity, whereas alanine substitutions^24, 25^, which has been used to reduce off-target activity, did not (Fig. S6c). The D892R substituted hfCas12Max was obviously more sensitive to mismatch, which suggests that D892R or G883R improved sgRNA binding specificity. Our data suggests that a semi-rational engineering strategy with arginine substitutions based on the EGFP-activated reporter system could be used as a general approach to improve the activity of CRISPR editing tools.

Through engineering, our Cas12i system has achieved high editing activity, high specificity and a broad PAM range, comparable to SpCas9, and better than other Cas12 systems. Given its smaller size, short crRNA guide, and self-processing features^4, 8, 10^, the Type V-I Cas12i system is suitable for *in vivo* multiplexed gene editing applications, including AAV^26^ or LNP^12, 13^.

In addition, we have confirmed that the Type V-I Cas12i system can be used in base editing applications. For base editor, the dCas12i system shows high A-to-G editing at A9-A11 sites even A19 of KLF locus, and C-to-T editing at A7-A10 sites, which is similar to the dCas12a system but is distinct from the dCas9/nCas9 system. Comparable to dCas12a, dCas12i-BE exhibited higher base editing activity at KLF4, PCSK9 and DYRK1A loci (Fig. 4c-d), suggesting it may have more potential as a base editor. This suggests that the dCas12i system is useful for broad genome engineering applications, including epigenome editing, genome activation, and chromatin imaging^1, 27-30^.

In summary, the Cas12i system described here, which has robust editing activity and high specificity, is a versatile platform for genome editing or base editing in mammalian cells and could be useful in the future for *in vivo* or *ex vivo* therapeutic applications.

Note that two groups have recently reported improved Cas12i2 using similar strategies in this study^31, 32^.

## Authors’ Contributions

H.Z., and Y.Z. jointly conceived and designed the project. H.Z., H.W., J.Z., W.Z., M.X., and X.K., performed experiments with plasmid construction, cell culture and FACS and data analysis. M.X., Z.W., Y.W., X.S., J.L., J.H., and N.Z. assisted with experiments. H.Y. supervised the whole project. H.Z., Y.Z., and H.Y. wrote and revised the manuscript.

## Material and methods

### Plasmid vector construction

Human codon-optimized *Cas12i, TadA8e* and human *APOBEC3A* genes were synthesized by the GenScript Co., Ltd., and cloned to generate pCAG_NLS-Cas12i-NLS_pA_pU6_BpiI_pCMV_mCherry_pA by Gibson Assembly. crRNA oligos were synthesized by HuaGene Co., Ltd., annealed and ligated into *Bpi*I site to produce the pCAG_NLS-Cas12i-NLS_pA_pU6_crRNA_pCMV_mCherry_pA.

### Cell culture, transfection and flow cytometry analysis

The mammalian cell lines used in this study were HEK293T and N2A. Cells were cultured in Dulbecco’s modified Eagle’s medium (DMEM) supplemented with 10% FBS, penicillin/streptomycin and GlutMAX. Transfections were performed using Polyetherimide (PEI). For variant screening, HEK293T cells were cultured in 24-well plates, and after 12 hours 2 μg of the plasmids (1 μg Cas12i of a mutant plasmid and 1 μg of the reporter plasmid) were transfected into these cells with 4 μL PEI. 48 hours after transfection, mCherry and EGFP fluorescence were analyzed using a Beckman CytoFlex flow-cytometer. For assay of mutations in target sites of endogenous genes, 1 μg plasmid with Cas12i targeting crRNA was transfected into HEK293T or N2A cells, which were then sorted using a BD FACS Aria III, BD LSRFortessa X-20 flow cytometer, 48 hours after transfection.

### Detection of gene editing frequency

Six thousand sorted cells were lysed in 20 μl of lysis buffer (Vazyme). Targeted sequence primers were synthesized and used in nested PCR amplification by Phanta Max Super-Fidelity DNA Polymerase (Vazyme). TIDE and targeted deep sequence analysis was used to determine indel frequencies. Sanger sequencing and EditR were used for quantification of base substitutions (A-to-G or C-to-T).

## Figure Legends

**Fig.1 hfCas12Max, an engineered natural variant xCas12i, mediated high-efficient and -specificity genome editing in mammalian cells. a**, Transfection of plasmids coding Cas12i and sgRNA activates EGFP. **b**, A natural variant xCas12i mediated EGFP activation efficiency determined by flow cytometry. **c**, Flow diagram for detection of genome editing efficiency by transfection of an all-in-one plasmid containing xCas12i and targeted gRNA into HEK293T cells, followed by FACS and NGS analysis. **d**, xCas12i mediated robust genome editing at the *Ttr* locus in N2a cells and *TTR* and *PCSK9* in HEK293T cells. **e-g**, Cas12Max exhibited significantly increased cleavage activity than xCas12i at various genomic or reporter plasmids target sites. **h**, Both Cas12Max and hfCas12Max exhibited a broader PAM recognition profile than other Cas proteins, including 5’-TN and 5’-TNN PAM. **i**, v6.3 reduced off-target at OT.1 and OT.2 sites and retained indel activity at *TTR*.*2*-ON targets, compared to v1.1-Cas12Max. **j**, TIDE analysis showed that v6.3-hfCas12Max retained comparable activity at TTR.2-ON targets and almost no at 6 OT sites, to Cas12Max. **k**, Comparison of indel activity from Cas12Max, hfCas12Max, LbCas12a, Ultra AsCas12a, SpCas9 and KKH-saCas9 at TTR locus. HfCas12Max retained the comparable activity of Cas12Max, and higher gene-editing efficiency than other Cas proteins. Each dot represents one of three repeats of single target site.

**Fig.2 dCas12Max-TadA8e and dCas12Max-hA3A.1 mediated high base editing activity. a**, Schematics of different versions of dCas12Max base editors. **b**, The A-to-G editing activity of TadA8e.1-dxCas12i-v2.2 was largely increased by combining v1.2 and v1.3 variant. **c**, Comparison of A-to-G editing frequency at the KLKF4 site from TadA8e.1-dxCas12i-v1.2, v2.2 and v4.3, ABE-dCas12Max (TadA8e-dxCas12i-v4.3) showed a high editing activity of 80%. **d**, Comparison of C-to-T editing frequency at the DYRK1A site from hA3A.1-dxCas12i, -v1.2 and v2.2, CBE-dCas12Max (hA3A.1-dxCas12i-v2.2) showed a high editing activity of 50%.

## Supplementary Figure Legends

**Supplementary Figure 1. Screen for functional Cas12i in HEK293T cells. a**, Five of ten natural Cas12i nuclease mediated EGFP-activated efficiency in HEK293T cells.

**Supplementary Figure 2. Identification and characterization of type V-I systems. a**, Nuclease domain organization of SpCas9, LbCas12a, and xCas12i. **b**, Optimal spacer length for xCas12i. **c**, PAM scope comparison of LbCas12a, and xCas12i. xCas12i exhibited a higher editing efficiency at 5’-TTN PAM than Cas12a. **d**, xCas12i mediated robust genome editing (up to 90%) at the *Ttr* locus in N2a cells and *TTR* and *PCSK9* in HEK293T cells.

**Supplementary Figure 3. Screen for engineered xCas12i mutants with high-efficiency editing activity. a**, Schematics of protein engineering strategy using an activatable EGFP reporter screening system. **b**, The relative editing frequencies of over 500 rationally engineered xCas12i mutants.

**Supplementary Figure 4. Other mutants mediated high-efficiency editing. a-b**, xCas12i mutant with N243R increased 1.2, 5, 20-fold activity at DMD.1, DMD.2 and DMD.3 locus. **c**, Both Cas12Max (xCas12i-N243R) and Cas12Max-E336R elevated EGFP-activated fluorescent at different PAM recognition sites.

**Supplementary Figure 5. Cas12Max induced off-target editing efficiency at sites with mismatches using the reporter system (a) and targeted deep sequence (b)**.

**Supplementary Figure 6. hfCas12Max mediates high-efficiency and -specificity editing. a**, Schematics of protein engineering strategy using EGFP activatable reporter screening system with on-targeted and off-targeted crRNA. **b**, Rational protein engineering screen of over 200 mutants for highly-fidelity Cas12Max.Four mutants show significantly decreased activity at both OT (off-target) sites and retains at ON.1 (on-target) site. **c**, Different versions of xCas12i mutants. **d**, v6.3-hfCas12Max reduced off-target at OT.1 and OT.2 sites and retained indel activity at *TTR*.*2*-ON targets, compared to v1.1-Cas12Max. **e**, v6.3-hfCas12Max exhibited comparable indel activity at DMD.1, DMD.2, and higher at DMD.3 locus, than v1.1-Cas12Max.

**Supplementary Figure 7. hfCas12Max exhibited the high editing activity. a**, Indel frenquencies from Cas12Max, hfCas12Max, LbCas12a, Ultra AsCas12a, SpCas9 and KKH-saCas9 at *TTR* locus.

**Supplementary Figure 8. hfCas12Max mediates high-efficient and -specific editing. a-b**, Off-target efficiency of hfCas12Max, LbCas12a, and UltraAsCas12a at in-silico predicted off-target sites, determined by targeted deep sequencing. Sequences of on-target and predicted off-target sites are shown, PAM sequences are in blue and mismatched bases are in red.

**Supplementary Figure 9. Conserved cleavage sites of Cas12i. a**, Sequence alignment of xCas12i, Cas12i1 and Cas12i2 shows that D650, D700, E875 and D1049 are conserved cleavage sites at RuvC domain. **b**, Introducing point mutations off D650A, E875A, and D1049A results in abolished activity of xCas12i.

**Supplementary Figure 10. Other strategies for high-efficiency ABE-dxCas12i. a**, TadA8e.1-dxCas12i-v1.2 and v1.3 exhibits significantly increased A-to-G editing activity among various variants at KLKF4 site of genome. **b**, Unchanged or even decreased editing activity from various dCas12-ABEs carrying different NLS at N-terminal. **c**, Increased A-to-G editing activity of TadA8e-dxCas12i-v4.3 by combining v2.2, changed-NLS linker and high-activity Tade8e. **d**, dxCas12i-ABE-N by TadA at the C-terminus of dCas12 slightly increased editing activity.

**Supplementary Figure 11. Comparison of editing frequencies induced by various dCas12-BEs at different genomic target sites. a**, Comparison of A-to-G editing frequencies induced by indicated TadA8e.1-dxCas12i, v1.2, v2.2, and TadA8e.1-dCas12a at *PCSK9* genomic locus.

**Supplementary Figure 1.**
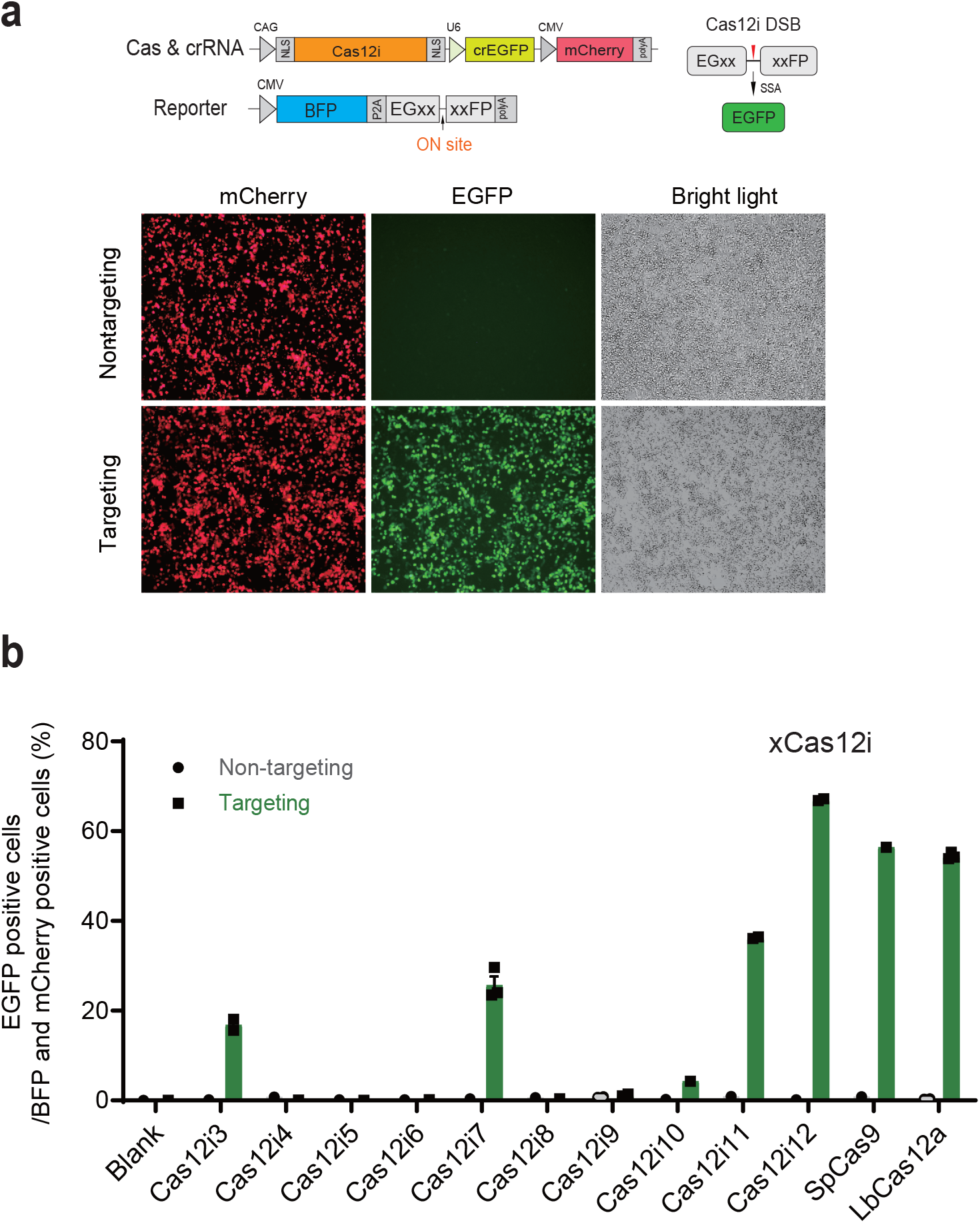
Screen for functional Cas12i in HEK293T cells. **a**, Transfection of plasmids coding Cas12i and crRNA mediated EGFP activation. **b**, Five of ten natural Cas12i nuclease mediated EGFP-activated efficiency in HEK293T cells.

**Supplementary Figure 2.**
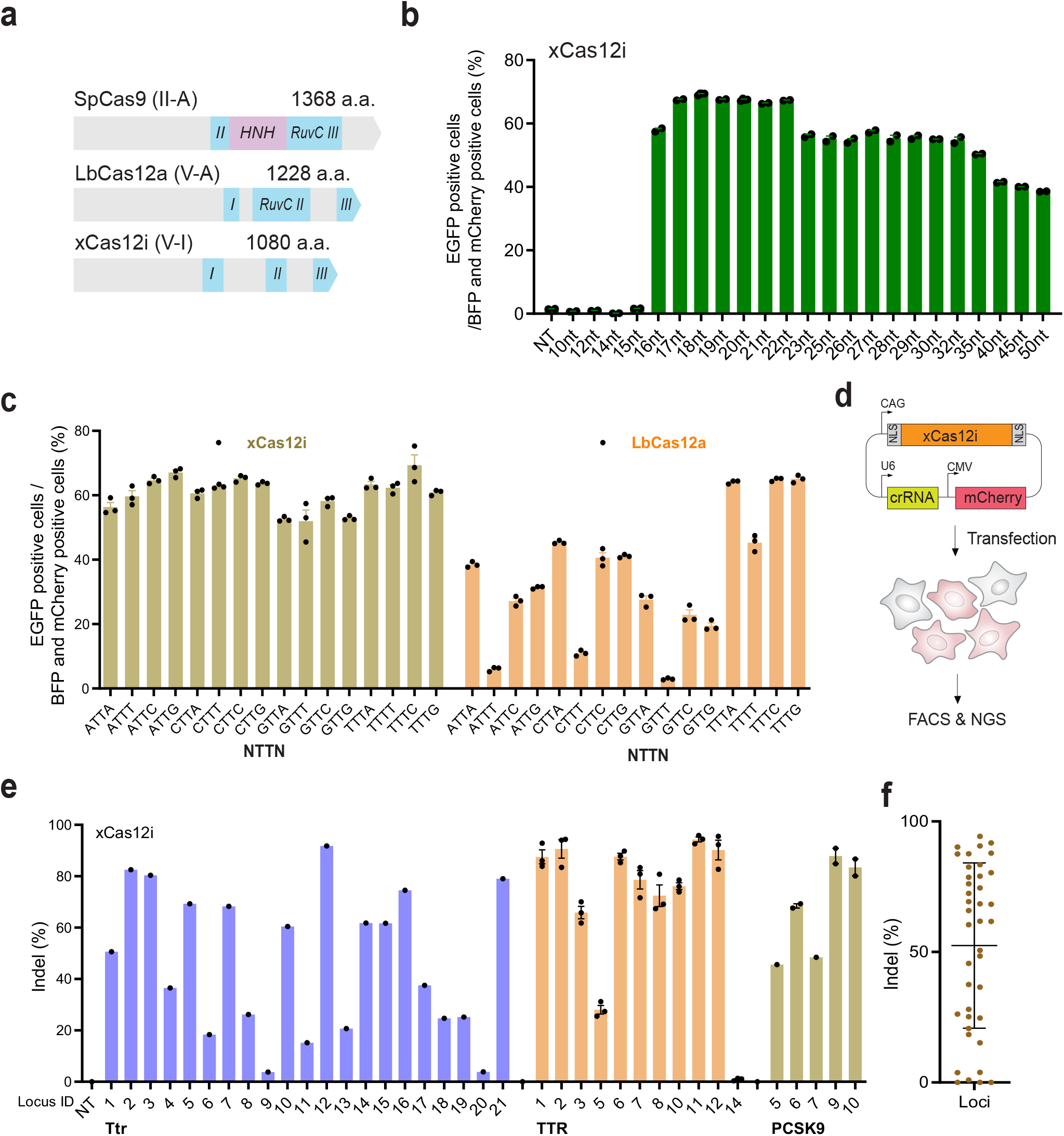
Identification and characterization of type V-I systems. **a**, Nuclease domain organization of SpCas9, LbCas12a, and xCas12i. **b**, Optimal spacer length for xCas12i. **c**, PAM scope comparison of LbCas12a, and xCas12i. xCas12i exhibited a higher editing efficiency at 5’-TTN PAM than Cas12a. **d**, Flow diagram for detection of genome editing efficiency by transfection of an all-in-one plasmid containing xCas12i and targeted gRNA into HEK293T cells, followed by FACS and NGS analysis. **e-f**, xCas12i mediated robust genome editing (up to 90%) at the Ttr locus in N2a cells and TTR and PCSK9 in HEK293T cells.

**Supplementary Figure 3.**
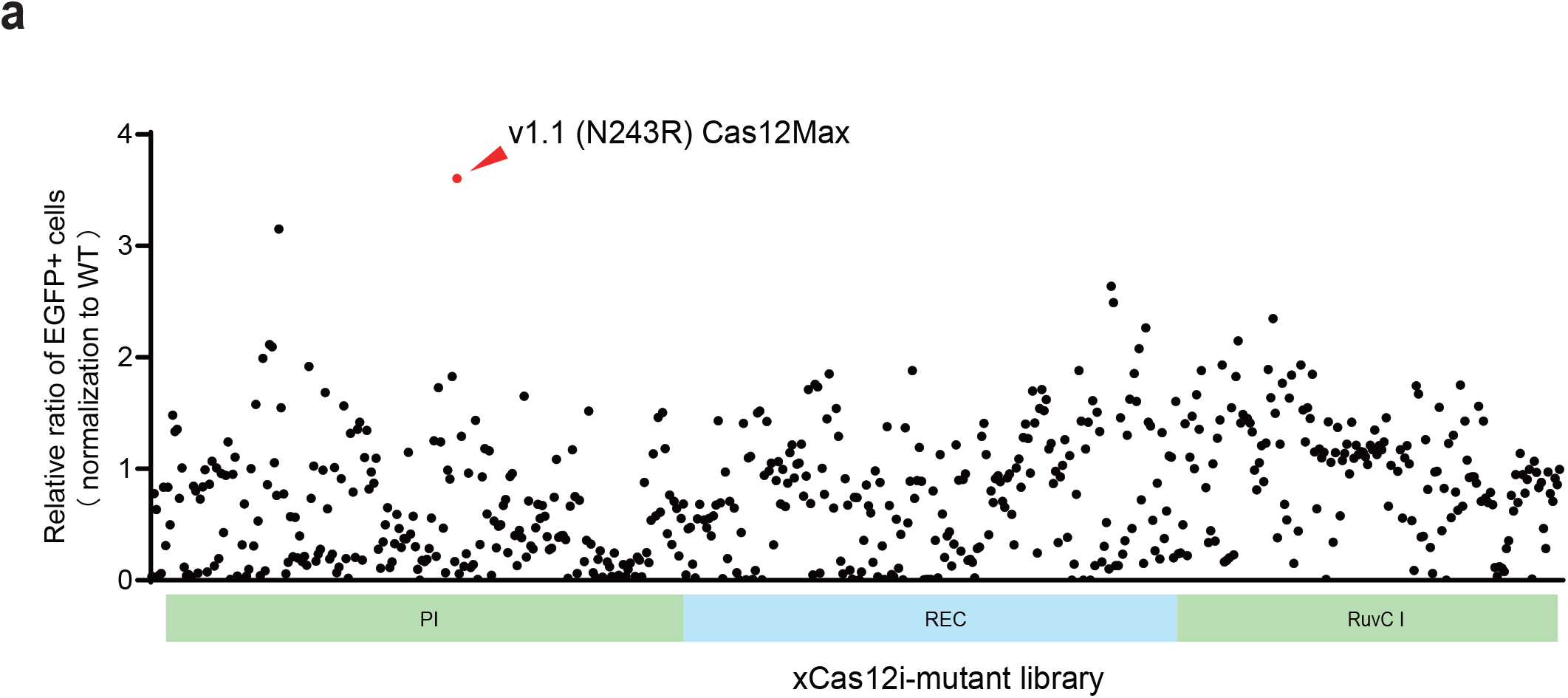
Screen for engineered xCas12i mutants with high-efficiency editing activity. **a**, The relative editing frequencies of over 500 rationally engineered xCas12i mutants.

**Supplementary Figure 4.**
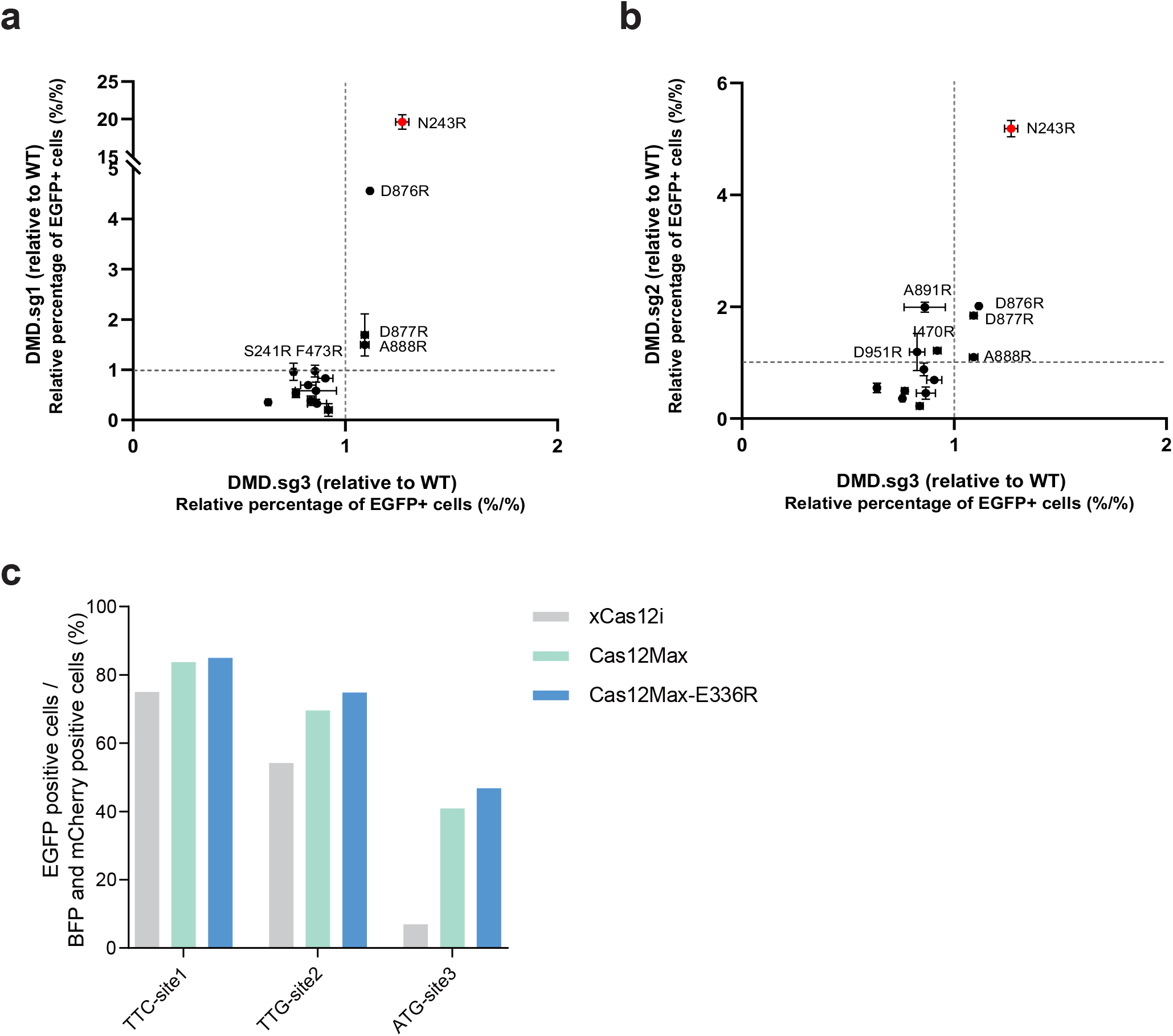
Other mutants mediated high-efficiency editing. **a-b**, xCas12i mutant with N243R increased 1.2, 5, 20-fold activity at DMD.1, DMD.2 and DMD.3 locus. **c**, Both Cas12-Max (xCas12i-N243R) and Cas12Max-E336R elevated EGFP-activated fluorescent at different PAM recognition sites.

**Supplementary Figure 5.**
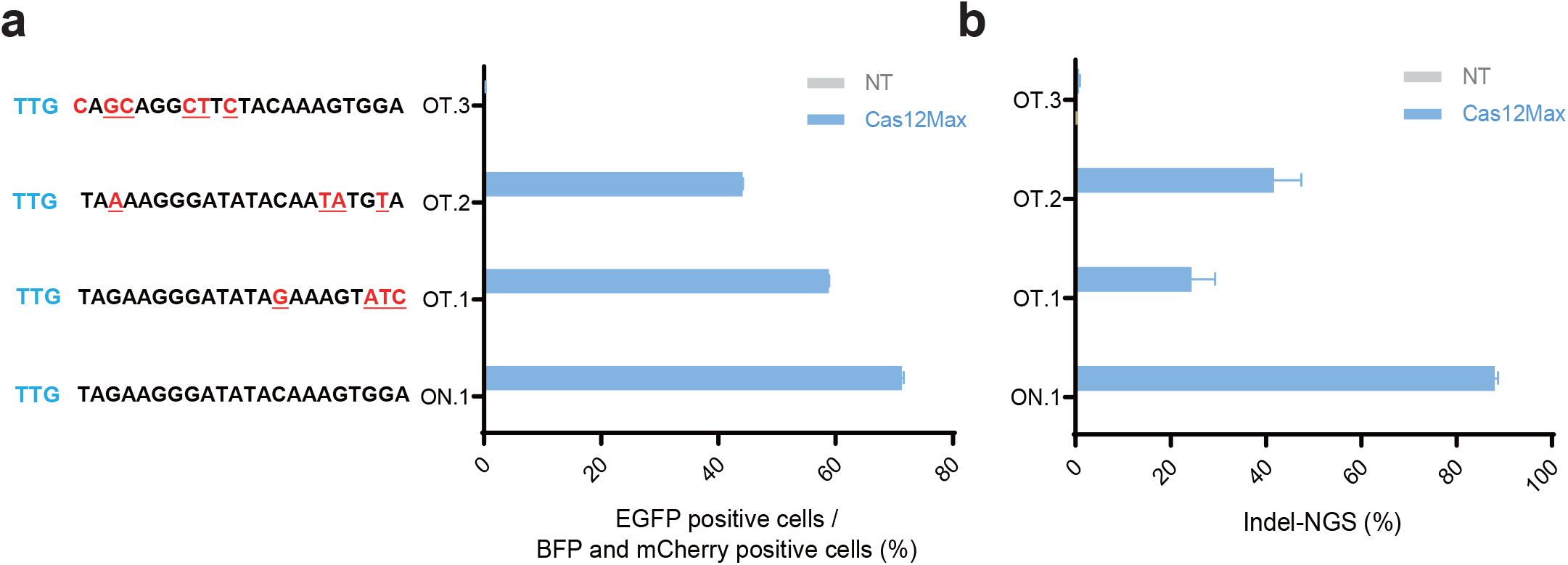
Cas12Max induced off-target editing efficiency at sites with mismatches using the reporter system (a) and targeted deep sequence (b).

**Supplementary Figure 6.**
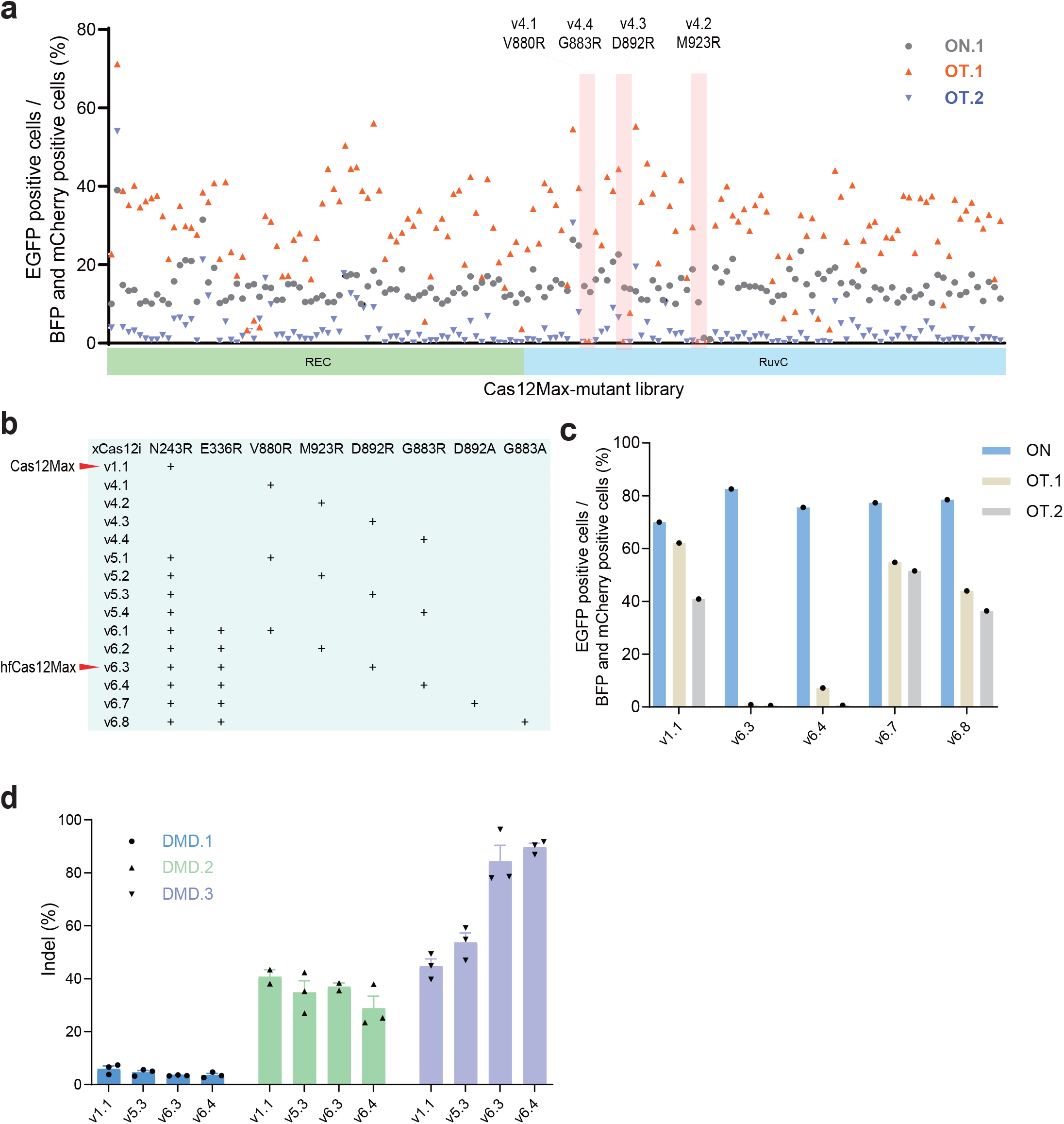
hfCas12Max mediates high-efficiency and -specificity editing. **a**, Rational protein engineering screen of over 200 mutants for highly-fidelity Cas12Max.Four mutants show significantly decreased activity at both OT (off-target) sites and retains at ON.1 (on-target) site. **b**, Different versions of xCas12i mutants. **c**, v6.3-hfCas12Max reduced off-target at OT.1 and OT.2 sites and retained indel activity at TTR-ON targets, compared to v1.1-Cas12-Max. **d**, v6.3-hfCas12Max exhibited comparable indel activity at DMD.1, DMD.2, and higher at DMD.3 locus, than v1.1-Cas12Max.

**Supplementary Figure 7.**
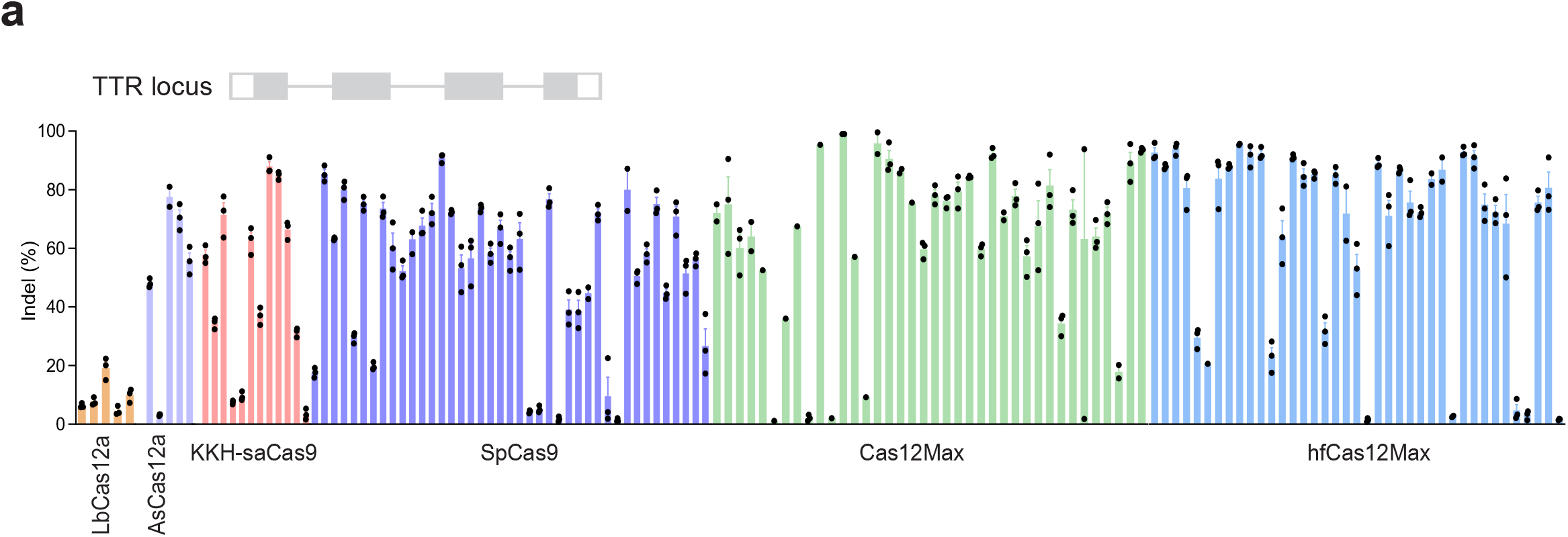
hfCas12Max exhibited the high editing activity. **a**, Indel frenquencies from Cas12Max, hfCas12Max, LbCas12a, Ultra AsCas12a, SpCas9 and KKH-saCas9 at TTR locus.

**Supplementary Figure 8.**
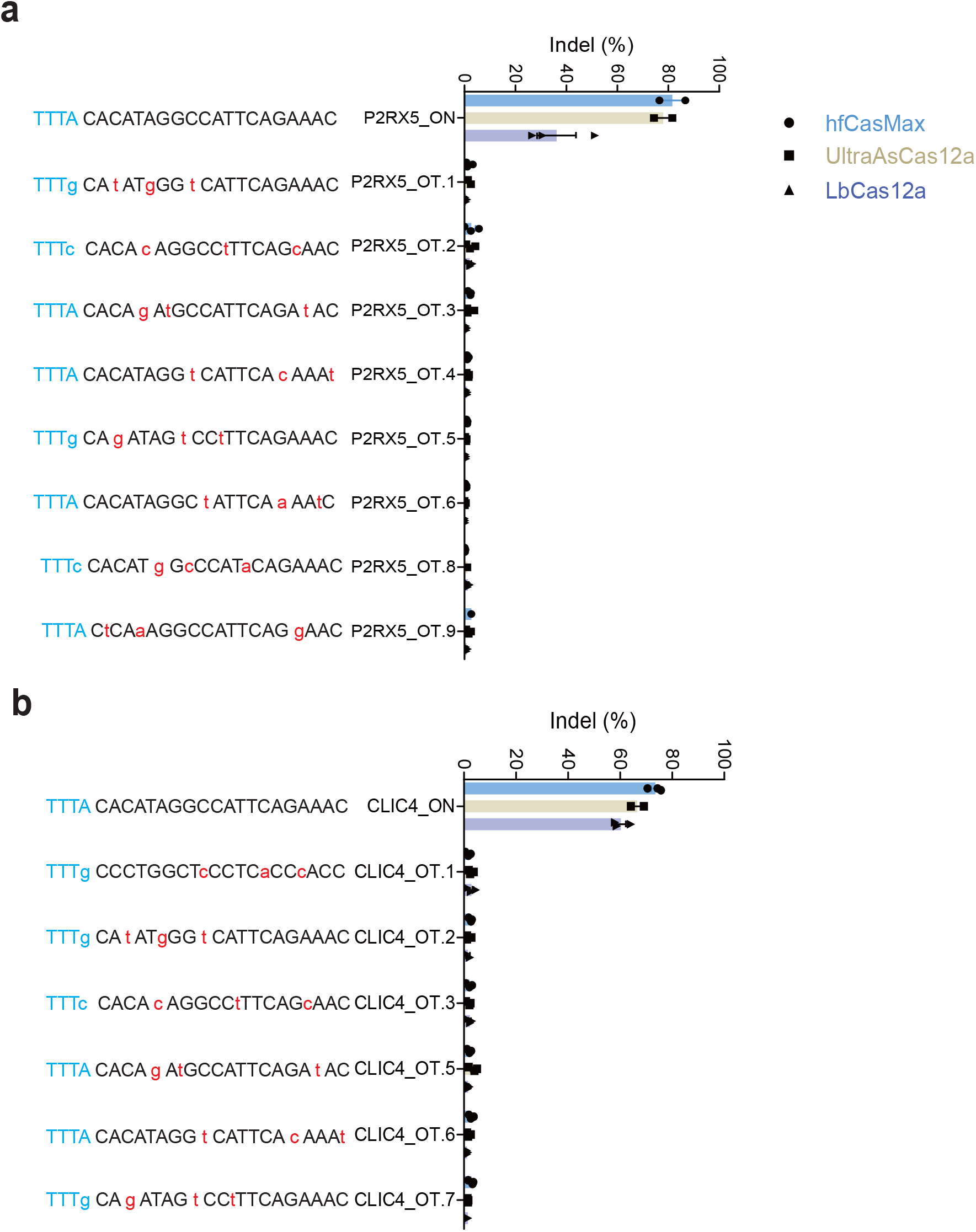
hfCas12Max mediates high-efficient and -specific editing. **a-b**, Off-target efficiency of hfCas12Max, LbCas12a, and UltraAsCas12a at *in-silico* predicted off-target sites, determined by targeted deep sequencing. Sequences of on-target and predicted off-target sites are shown, PAM sequences are in blue and mismatched bases are in red.

**Supplementary Figure 9.**
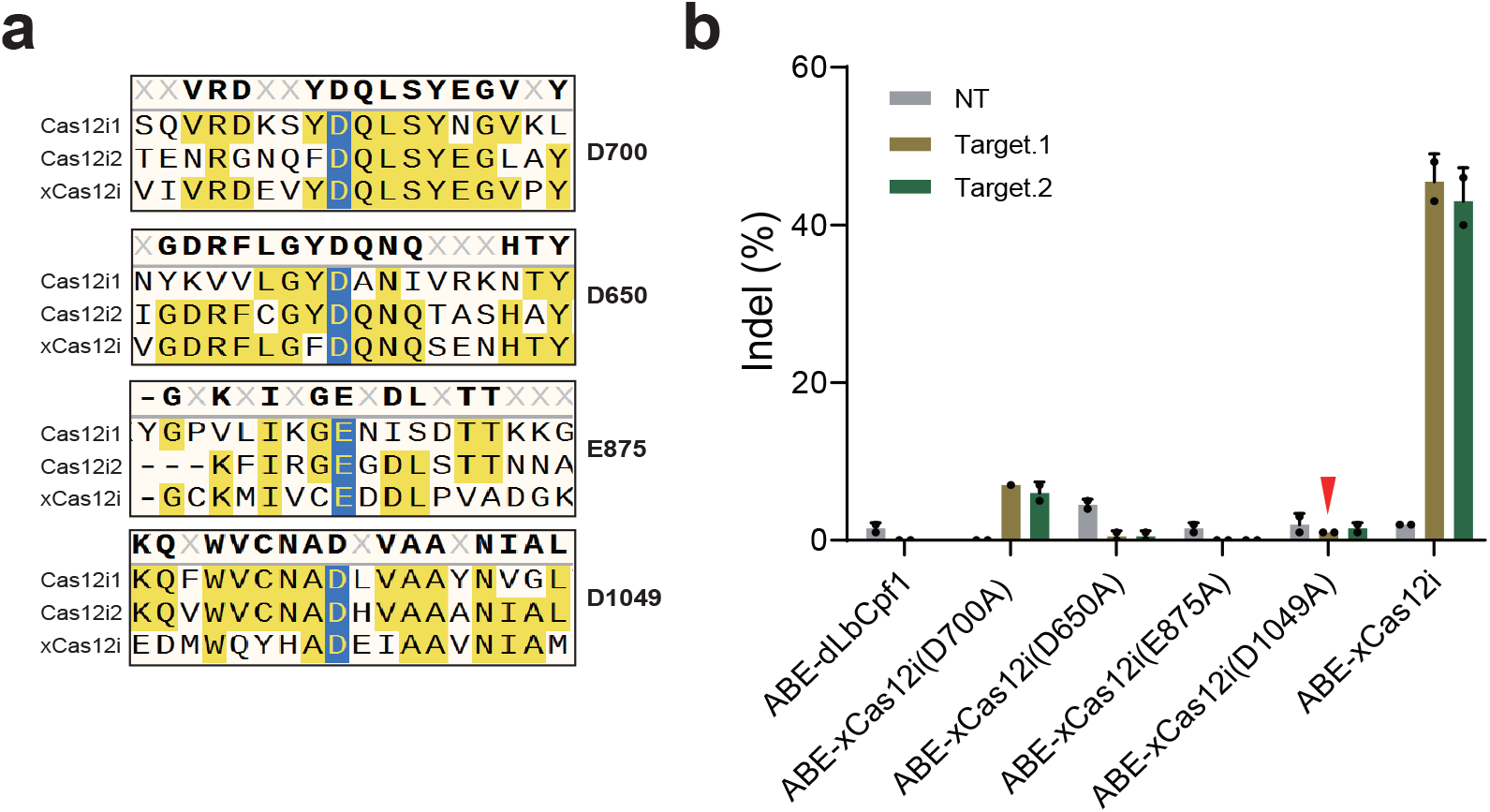
Conserved cleavage sites of Cas12i. **a**, Sequence alignment of xCas12i, Cas12i1 and Cas12i2 shows that D650, D700, E875 and D1049 are conserved cleavage sites at RuvC domain. **b**, Introducing point mutations off D650A, E875A, and D1049A results in abolished activity of xCas12i.

**Supplementary Figure 10.**
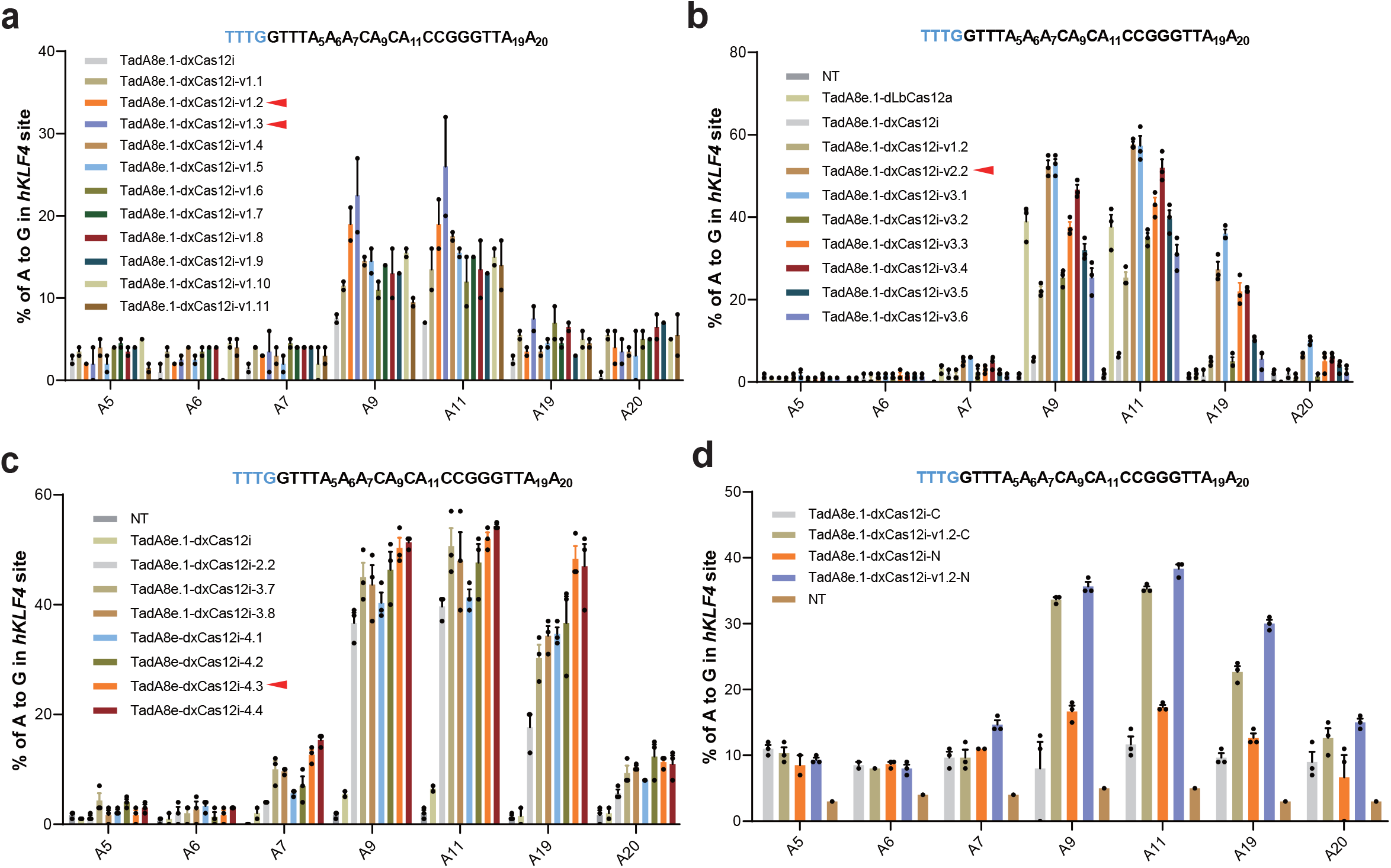
Other strategies for high-efficiency ABE-dxCas12i. **a**, TadA8e.1-dxCas12i-v1.2 and v1.3 exhibits significantly increased A-to-G editing activity among various variants at KLKF4 site of genome. **b**, Unchanged or even decreased editing activity from various dCas12-ABEs carrying different NLS at N-terminal. **c**, Increased A-to-G editing activity of TadA8e-dxCas12i-v4.3 by combining v2.2, changed-NLS linker and high-activity Tade8e. **d**, dxCas12i-ABE-N by TadA at the C-terminus of dCas12 slightly increased editing activity.

**Supplementary Figure 11.**
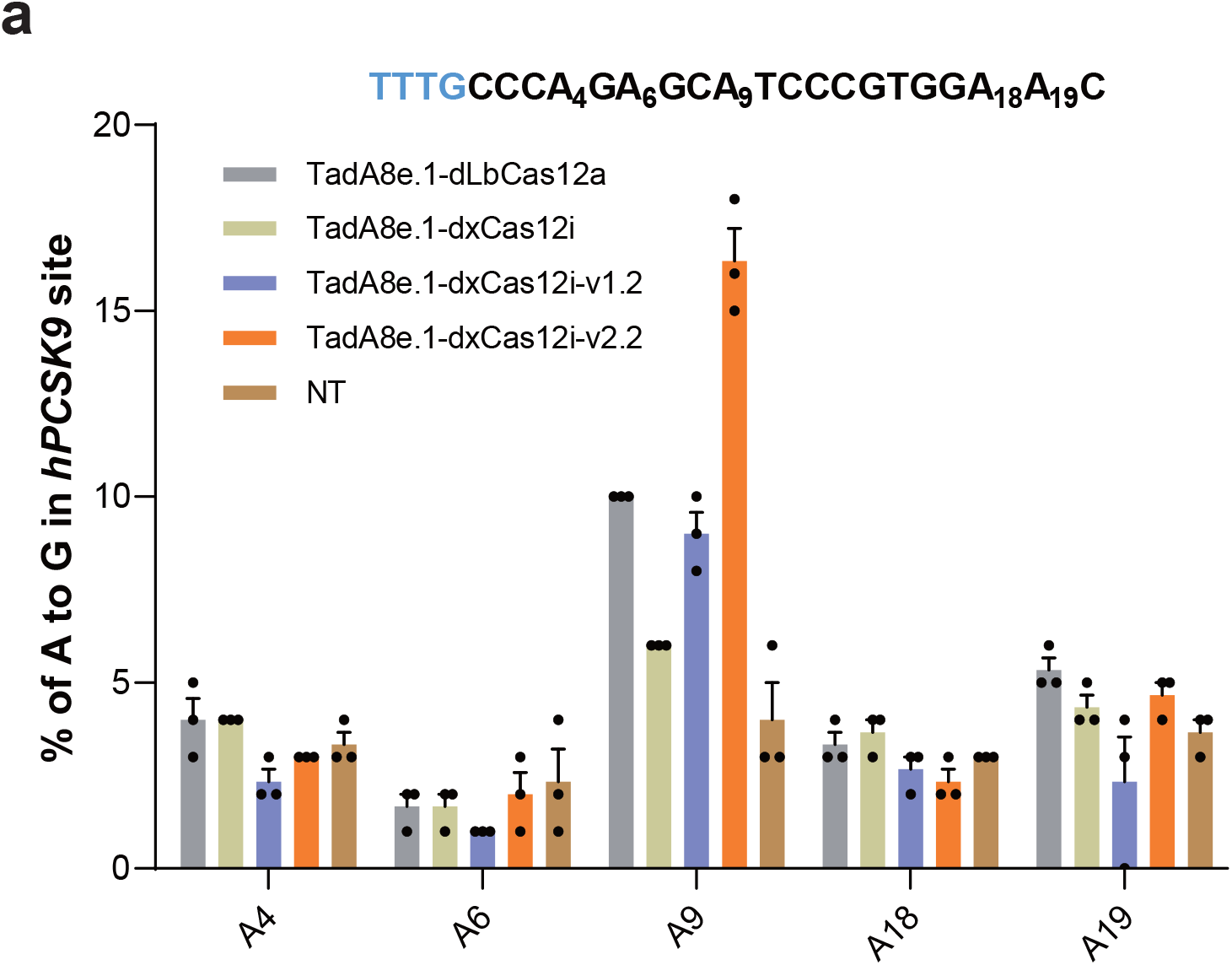
Comparison of editing frequencies induced by various dCas12-BEs at different genomic target sites. **a**, Comparison of A-to-G editing frequencies induced by indicated TadA8e.1-dxCas12i, v1.2, v2.2, and TadA8e.1-dCas12a at PCSK9 genomic locus.

## References

1. Anzalone, A.V., Koblan, L.W. & Liu, D.R. Genome editing with CRISPR-Cas nucleases, base editors, transposases and prime editors. Nat Biotechnol 38, 824–844 (2020).

2. Doudna, J.A. The promise and challenge of therapeutic genome editing. Nature 578, 229–236 (2020).

3. Makarova, K.S. et al. Evolutionary classification of CRISPR-Cas systems: a burst of class 2 and derived variants. Nat Rev Microbiol 18, 67–83 (2020).

4. Yan, W.X. et al. Functionally diverse type V CRISPR-Cas systems. Science 363, 88–91 (2019).

5. Kleinstiver, B.P. et al. Genome-wide specificities of CRISPR-Cas Cpf1 nucleases in human cells. Nature Biotechnology 34, 869-+ (2016).

6. Cong, L. et al. Multiplex Genome Engineering Using CRISPR/Cas Systems. Science 339, 819–823 (2013).

7. Zetsche, B. et al. Cpf1 is a single RNA-guided endonuclease of a class 2 CRISPR-Cas system. Cell 163, 759–771 (2015).

8. Zhang, B. et al. Mechanistic insights into the R-loop formation and cleavage in CRISPR-Cas12i1. Nature communications 12, 3476 (2021).

9. Zhang, H., Li, Z., Xiao, R. & Chang, L. Mechanisms for target recognition and cleavage by the Cas12i RNA-guided endonuclease. Nat Struct Mol Biol 27, 1069–1076 (2020).

10. Huang, X. et al. Structural basis for two metal-ion catalysis of DNA cleavage by Cas12i2. Nature communications 11, 5241 (2020).

11. Yang, Y. et al. Highly Efficient and Rapid Detection of the Cleavage Activity of Cas9/gRNA via a Fluorescent Reporter. Appl Biochem Biotechnol 180, 655–667 (2016).

12. Gillmore, J.D. et al. CRISPR-Cas9 In Vivo Gene Editing for Transthyretin Amyloidosis. The New England journal of medicine 385, 493–502 (2021).

13. Musunuru, K. et al. In vivo CRISPR base editing of PCSK9 durably lowers cholesterol in primates. Nature 593, 429–434 (2021).

14. Strecker, J. et al. Engineering of CRISPR-Cas12b for human genome editing. Nature communications 10, 212 (2019).

15. Kleinstiver, B.P. et al. Engineered CRISPR-Cas12a variants with increased activities and improved targeting ranges for gene, epigenetic and base editing. Nat Biotechnol 37, 276–282 (2019).

16. Xu, X. et al. Engineered miniature CRISPR-Cas system for mammalian genome regulation and editing. Mol Cell 81, 4333–4345 e4334 (2021).

17. Zhang, L. et al. AsCas12a ultra nuclease facilitates the rapid generation of therapeutic cell medicines. Nature communications 12, 3908 (2021).

18. Bae, S., Park, J. & Kim, J.S. Cas-OFFinder: a fast and versatile algorithm that searches for potential off-target sites of Cas9 RNA-guided endonucleases. Bioinformatics 30, 1473–1475 (2014).

19. Yuen, C.T.L. et al. High-fidelity KKH variant of Staphylococcus aureus Cas9 nucleases with improved base mismatch discrimination. Nucleic Acids Res 50, 1650–1660 (2022).

20. Kim, D.Y. et al. Efficient CRISPR editing with a hypercompact Cas12f1 and engineered guide RNAs delivered by adeno-associated virus. Nat Biotechnol 40, 94–102 (2022).

21. Wang, X. et al. Cas12a Base Editors Induce Efficient and Specific Editing with Low DNA Damage Response. Cell Rep 31, 107723 (2020).

22. Richter, M.F. et al. Phage-assisted evolution of an adenine base editor with improved Cas domain compatibility and activity. Nat Biotechnol 38, 883–891 (2020).

23. Li, X. et al. Base editing with a Cpf1-cytidine deaminase fusion. Nat Biotechnol 36, 324–327 (2018).

24. Bravo, J.P.K. et al. Structural basis for mismatch surveillance by CRISPR-Cas9. Nature 603, 343–347 (2022).

25. Kleinstiver, B.P. et al. High-fidelity CRISPR-Cas9 nucleases with no detectable genome-wide off-target effects. Nature 529, 490–495 (2016).

26. Wang, D., Zhang, F. & Gao, G. CRISPR-Based Therapeutic Genome Editing: Strategies and In Vivo Delivery by AAV Vectors. Cell 181, 136–150 (2020).

27. Wang, H. et al. CRISPR-Mediated Programmable 3D Genome Positioning and Nuclear Organization. Cell 175, 1405–1417 e1414 (2018).

28. Konermann, S. et al. Genome-scale transcriptional activation by an engineered CRISPR-Cas9 complex. Nature 517, 583–588 (2015).

29. Nakamura, M., Gao, Y., Dominguez, A.A. & Qi, L.S. CRISPR technologies for precise epigenome editing. Nat Cell Biol 23, 11–22 (2021).

30. Fellmann, C., Gowen, B.G., Lin, P.C., Doudna, J.A. & Corn, J.E. Cornerstones of CRISPR-Cas in drug discovery and therapy. Nat Rev Drug Discov 16, 89–100 (2017).

31. Chen, Y. et al. Synergistic engineering of CRISPR-Cas nucleases enables robust mammalian genome editing. Innovation (Camb) 3, 100264 (2022).

32. McGaw, C. et al. Engineered Cas12i2 is a versatile high-efficiency platform for therapeutic genome editing. Nature communications 13, 2833 (2022).

